# Task-specific neural mechanisms underlie biases in human orientation perception

**DOI:** 10.64898/2026.07.12.738112

**Authors:** Richard Leadbeater, Timothy Ledgeway, Paul McGraw

**Affiliations:** Department of Psychology, University of Nottingham, Nottingham, United Kingdom

## Abstract

Prior experience shapes both visual perception and its underlying neural circuits. This is exemplified by the *oblique effect* – a strong perceptual advantage for cardinal (horizontal/vertical) over oblique orientations – which reflects how the brain adapts to statistical regularities in the natural environment. It remains unclear whether such adaptations are generalised across visual cortex or are specific to circuits supporting different perceptual judgements. To investigate, we examined human performance in contrast detection and orientation discrimination, using identical stimuli for a range of spatial frequencies, paired with a biologically-inspired model of visual orientation processing. Behaviourally, a robust oblique effect emerged for orientation discrimination but was found only at higher frequencies for contrast detection. The model explained detection changes via an increased pooled response from cardinal-tuned neurons alongside spatial frequency-dependent narrowing of orientation bandwidths, consistent with known properties of cortical V1 neurons. However, the discrimination oblique effect required a different constraint, narrower orientation tuning for cardinal versus oblique neurons. No single model captured both effects simultaneously, suggesting that the oblique effect results from task-specific mechanisms. More broadly, these findings demonstrate how, rather than relying on a fixed strategy, the brain employs flexible computational strategies to optimise sensory encoding for specific tasks.

**Author Summary:** Everyday scenes are dominated by horizontal and vertical contours, such as horizons, buildings, and other natural or man-made structures. The human visual system appears tuned to this regularity: people judge horizontal and vertical orientations more accurately than oblique ones. This bias is thought to arise from neural mechanisms that encode orientation, and we investigated whether this reflects a fixed property of orientation coding, or flexible adaptation to task demands. Participants performed a *detection* task, in which they reported the presence or absence of faint oriented patterns, and a *discrimination* task, in which they judged small changes in orientation. We built a computational model based on known response properties of orientation-selective neurons in visual cortex, to test which neural adaptations best-explained performance on each task. Participants exhibited advantages for horizontal and vertical orientations which differed between the two tasks. Critically, our model revealed that this bias could not be explained by a single, general-purpose neural adaptation applied uniformly across both tasks. Instead, our data suggest that the visual system contains distinct orientation-coding biases that are engaged in a task-dependent manner. Consequently, sensory processing is shaped not only by regularities of the environment, but also by the observer’s behavioural goals.

## Introduction

The oblique effect is characterised by poorer visual performance for stimuli presented at oblique orientations, compared to cardinal orientations, on a variety of psychophysical tasks [1,2], and is thought to reflect an adaptation to the prominence of cardinal orientations in the visual environment [3–5]. Extensive psychophysical and neurophysiological work has informed hypotheses regarding the neural mechanisms of the oblique effect, but these cannot be tested experimentally owing to an inherent lack of control in experimental settings – experimenters cannot simply rewire visual cortex and measure the consequent changes in visual function. Furthermore, it is unknown if orientation biases on different visual tasks are governed by a shared neuronal adaptation to environmental statistics, or by multiple, task-specific adaptations.

The oblique effect for contrast detection (i.e., contrast sensitivity) is absent at low spatial frequencies (coarse spatial scales) but present at higher spatial frequencies, appearing at approximately >8 c/° in central vision [6,7]. Contrast sensitivity can be predicted by the amplitude of pooled activity recorded from large populations of neurons in early visual cortex [8–10]. Thus, anisotropic contrast sensitivity is often attributed to an overrepresentation of neurons selective for cardinal orientations [11], such that cardinal orientations elicit larger responses than oblique orientations [12,13]. Indeed, functional magnetic resonance imaging (fMRI) in humans [14] and optical imaging in macaques [15] reveal that a greater extent of primary visual cortex (V1) – the first visual area in higher mammals to exhibit strong orientation preferences [16] – is selective for cardinal than oblique orientations. However, this cardinal bias is also recorded in response to stimuli at lower spatial frequencies where, counterintuitively, the behavioural oblique effect is not found for contrast detection [14,15]. This is investigated using a computational modelling approach in the current study.

Unlike contrast sensitivity, the ability to discriminate small changes in the orientation of a stimulus is thought to depend on the slopes of orientation-tuning curves [17]. For instance, a small change in orientation will elicit a larger change in response for a neuron with a narrow tuning curve (steep slope) relative to a neuron with a broad tuning curve (shallow slope). Thus, performance on orientation-discrimination tasks is considered to reflect the precision of orientation encoding in the visual system. It follows that the oblique effect for orientation discrimination [18,19] could potentially be explained by narrower orientation bandwidths for neurons preferring cardinal than oblique orientations [20,21].

In apparent contradiction to the proposed mechanisms of the oblique effect in detection and discrimination tasks, single-cell recordings tend to demonstrate little to no systematic modulation of the number of neurons or orientation bandwidth with respect to the preferred orientation of macaque V1 or V2 neurons [22,23]. This lends support to the idea that distinct orientation anisotropies may occur in separate neural circuits supporting particular visual functions, rather than existing as a general property of early visual cortex. Task-specific adaptations in visual function are well-documented in perceptual learning studies. For example, an increased population response in cat striate cortex was found after training on a detection task [24,25] compared to a slight decrease after training on an orientation-discrimination task [26]. Similarly, orientation-discrimination training in primates is associated with narrowed orientation bandwidths near the trained orientation in extrastriate visual area V4 [27,28], whereas training on a contrast-discrimination task did not alter V4 orientation bandwidths [29]. One way to examine whether task-specific mechanisms underlie orientation biases on different visual tasks is to run model simulations on analogous computational tasks, and evaluate whether a single, shared model can explain the behavioural data on both tasks.

Previous modelling work reports that the oblique effect for motion-direction discrimination (i.e. a selective disadvantage for discriminating oblique movements) can be simulated by a neuronal population with anisotropic motion-direction bandwidths, and a concurrent overrepresentation of neurons selective to cardinal directions [30]. However, no systematic comparisons were made to assess how different levels of each orientation anisotropy influenced model performance. One study that implemented a comprehensive range of parameter values found that the oblique effect for estimates of motion direction was best explained by narrowed cardinal bandwidths [31]. However, their behavioural data were likely influenced by cognitive factors associated with subjective estimation and reproduction tasks [32,33]. Further simulations are therefore needed to clarify the mechanisms underlying biases in the encoding of motion-direction, and orientation, in human vision.

To date, there are no previous computational models of the oblique effect for contrast detection, nor investigations into whether a shared mechanism can explain orientation biases associated with different types of orientation judgement. To address this, we implemented a neurophysiologically-inspired model of orientation encoding, comprising populations of visual neurons with response parameters informed by neurophysiological work in macaque V1, to simulate performance at cardinal and oblique orientations on contrast-detection and orientation-discrimination tasks. Model performance was simulated in the presence of various orientation anisotropies (number of neurons, orientation bandwidth, response gain, response variability) which were adjusted to best-fit human psychophysical performance on analogous behavioural tasks. To investigate how orientation information may be encoded and then subsequently decoded in human vision, model performance was compared for different pooling and “read-out” algorithms. Our results are interpreted on the assumption that if an iteration of the model fails to capture patterns in the behavioural data, then such a model is unlikely to reflect the neural mechanisms underlying human visual performance. If a shared model can replicate the behavioural oblique effect on both tasks, this would indicate that orientation is encoded by a general mechanism. Conversely, the failure of a shared model provides evidence that the brain employs flexible computational strategies to optimise sensory encoding on specific tasks.

## Methods

### Participants

Eight participants took part in both the contrast-detection and orientation-discrimination experiments. All but three participants (P1, P2, and P3) were naïve to the aims of the study. The mean (SD) participant age was 34.1 (12.7) years. All had normal or corrected-to-normal vision and gave written, informed consent. Experimental procedures were approved by the Ethics Committee of the School of Psychology, University of Nottingham (*Approval Number: S1257R*). The recruitment period ran from 01/01/2021 to 15/12/2022.

### Apparatus and Stimuli

Visual stimuli were presented on a 20-inch CRT monitor (*iiyama Vision Master Pro 514*) with a refresh rate of 60 Hz, display resolution of 1152 x 870 pixels, and a maximum luminance of 52.24 cd/m^2^. Monitor luminance output was carefully linearised through photometric gamma correction (*Minolta LS110 Luminance Meter*). Precise control over luminance resolution was achieved by a widely used dithering algorithm, the “noisy-bit” method [45]. Testing was conducted in a darkened, quiet room and the viewing distance was fixed at 126 cm by means of a chinrest, such that the angular extent of each pixel was 0.0156 °. Stimulus generation and response collection were achieved using PsychoPy [46] and custom-written Python code.

Stimuli were Gabors, composed of sinusoidal gratings presented within a spatially two-dimensional Gaussian window with a space-constant *σ*^2^ of 1.33 °, truncated at ± 4 °. The standard orientation (either 0, 45, 90, or 135 °) and spatial frequency (either 2, 8, or 16 c/°) conditions set the corresponding parameters of the standard stimulus, which were constant throughout each run of trials. At 16 c/°, the sine wave was sampled at four pixels per cycle, above the Nyquist limit for accurate representation of the stimulus. The minimum, mean and maximum luminance values of the sine wave were always sampled. On each stimulus presentation, the phase was randomised (by shifting the sinusoid ± half of a cycle) to minimise the influence of luminance artifacts and prevent local adaptation across trials.

### Procedure

#### i. Staircase

Thresholds were determined using a temporal, two-alternative, forced-choice task (2AFC) with a 3-down 1-up, adaptive staircase procedure that tracked the 79.4 % correct performance level. Increments and decrements in the staircase were set on a log-scale for contrast detection and a linear scale for orientation discrimination. Detection thresholds were calculated as the geometric mean, and discrimination thresholds as the arithmetic mean, of the last five reversals in each run of trials. Each run ended after ten reversals, and a minimum of four runs was completed for each experimental condition. All runs were completed in a randomised order. Thresholds were calculated for each combination of orientation and spatial frequency conditions, except for two participants (P7 and P8) who were unavailable for data collection at 8 c/° for the experiments.

#### ii. Trials

On each trial (Figure 1), stimuli were presented in one of two temporal intervals, each with a duration of 300 ms, separated by a homogeneous “grey” blank field for 500 ms. Intervals were marked by distinct audible tones. A central fixation spot was presented at the end of each trial until a valid keyboard response was collected. A response triggered the initiation of the next trial after a 500 ms delay. Participants made judgements using the “F” and “J” keys to refer to the first and second intervals, respectively. No response feedback concerning the correctness of the participant’s decision was provided.

**Figure 1.**
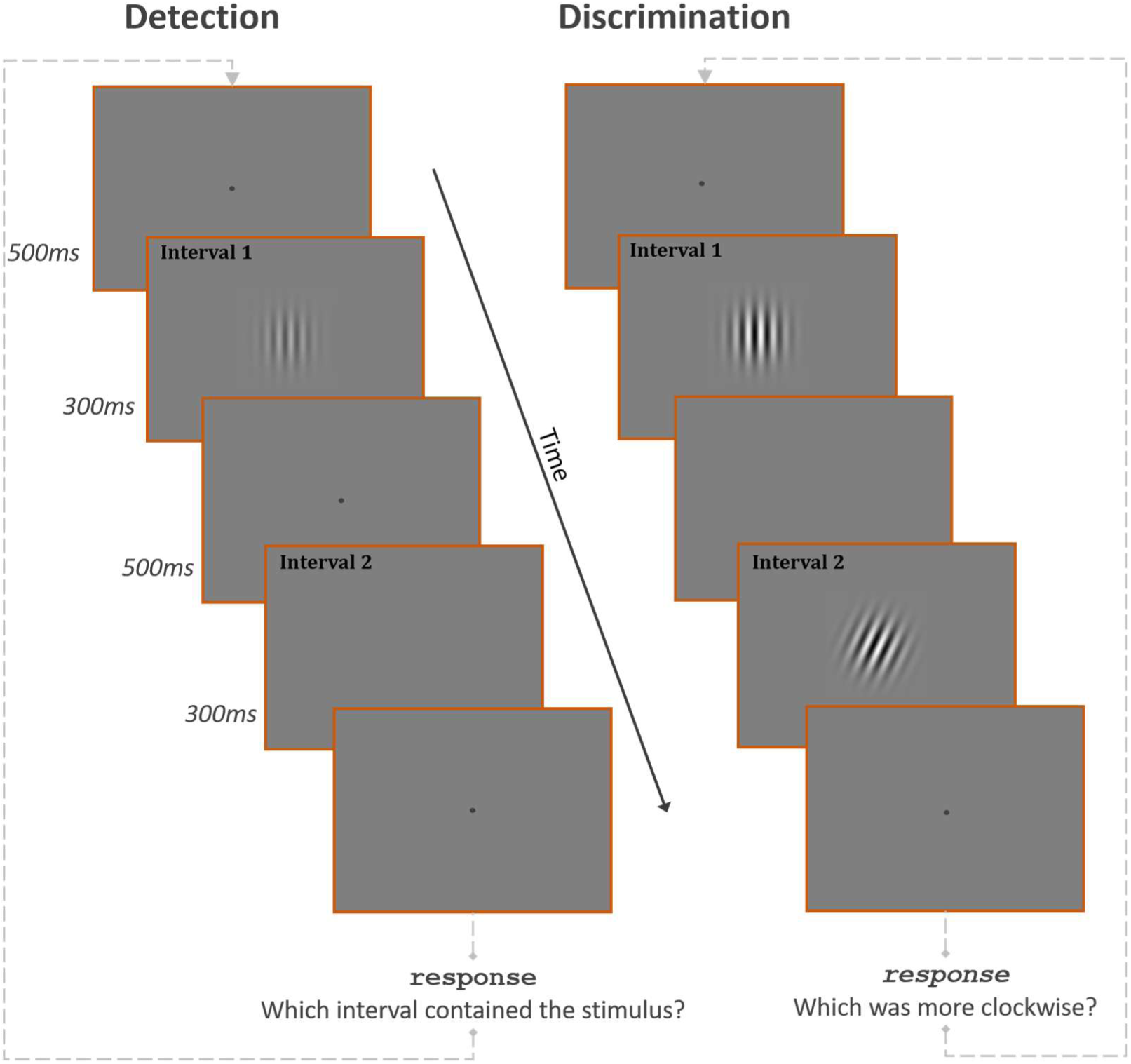
The time-course of the procedure for contrast-detection (*left*) and orientation-discrimination (*right*) tasks. Intervals 1 and 2 were signalled by distinct audible tones. In the detection task, participants judged whether the stimulus appeared in the first or second interval. In the discrimination task, participants judged whether the more clockwise grating appeared in the first or second interval.

#### iii. Experiment 1 – Contrast detection

The standard stimulus was presented in one of two temporal intervals, which was randomised on each trial. No stimulus was presented in the other interval. Participants judged which interval contained the standard stimulus. On each trial the Michelson contrast (%) of the standard stimulus was set according to the adaptive staircase procedure.

#### iv. Experiment 2 – Orientation discrimination

The standard stimulus and a comparison stimulus were presented sequentially in one of two temporal intervals, which was randomised on each trial. The orientation of the comparison stimulus was set relative to the standard orientation, and participants judged which interval contained the stimulus that was “more clockwise”. On each trial, the angular difference between the standard and comparison orientations was set according to the staircase, and the sign of the angular difference (i.e., direction of rotation) was randomised.

To ensure that stimulus visibility was equated across all tested conditions, stimulus contrast was conventionally set to a fixed multiple of each participant’s contrast threshold in the corresponding orientation and spatial frequency conditions. The fixed multiple *m* for each participant was calculated by dividing the maximum available contrast (nominally 100 %) by their highest detection threshold across all conditions (Table 1). Detection thresholds were not known *a priori*, therefore *m* was defined separately for each participant to ensure that stimuli were presented at the highest feasible suprathreshold level without exceeding the 100 % contrast ceiling in any condition.

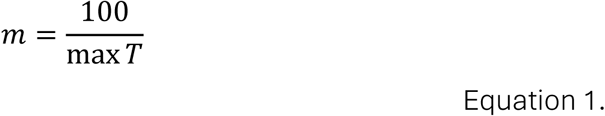

**Table 1.** The multiple *m* of the contrast threshold used for each participant.

|  | P1 | P2 | P3 | P4 | P5 | P6 | P7 | P8 |
| --- | --- | --- | --- | --- | --- | --- | --- | --- |
| m | 24.6 | 6.42 | 6.75 | 3.69 | 25.7 | 4.92 | 16.4 | 15.8 |

### Oblique efect index

The size of oblique effect was quantified by the “oblique effect index” (OEI), a ratio measure which controls for differences in overall performance between individuals and experimental conditions [19]:

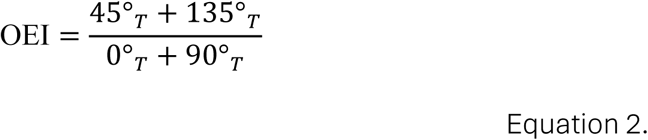

Where subscript *T* denotes a threshold value for a given orientation. Values above one indicates the presence of an oblique effect and quantify its magnitude (i.e., an OEI score of 2 shows that oblique thresholds are twice as high as cardinal thresholds).

### Model

#### i. Orientation tuning

To capture the behavioural data of Experiments 1 and 2, we implemented a neurophysiologically-inspired computational model of orientation encoding. Model neurons had a von Mises sensitivity profile, where the sensitivity of the *i*th neuron to orientation *θ* is:

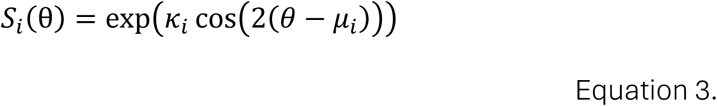

Where *μ*_i_ is the preferred orientation and *κ*_i_ is a width parameter of the *i*th neuron. Each neuron’s sensitivity profile was subsequently rescaled such that its minimum and maximum values were mapped to 0 and 1, respectively. The orientation bandwidth ℎ_i_ (half-width half-max, HWHM) in radians is given by:

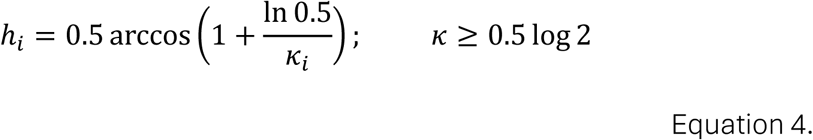

Tuned responses were modulated by stimulus contrast according to the conventional Naka-Rushton function [47,48]:

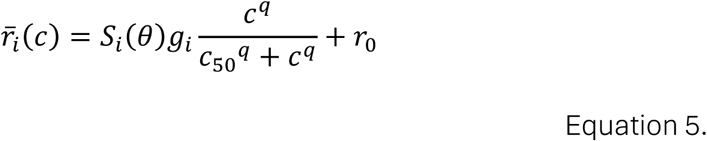

Where *r*_i_(*c*) is the tuned response of the *i*th neuron at Michelson contrast *c*. The semi-saturation constant (*c*_sO_ = 24), exponent (*q* = 2), and spontaneous firing rate (*r*_O_ = 3 sp/s) were fixed according to average measurements from macaque V1 [47,49]. The response gain *g*_i_ determined the maximum firing rate of each neuron, with a default maximum firing rate of 60 sp/s.

#### ii. Internal noise

To simulate the well-known variability of firing rates in visual cortical neurons [50], noise was added to the tuned response on a trial-to-trial basis. The noisy firing rate of the *i*th neuron is:

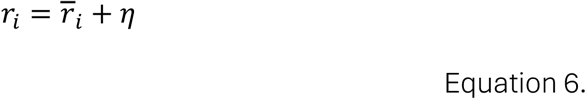

Where *η* is an independent random variable drawn from a Gaussian distribution centred on zero:

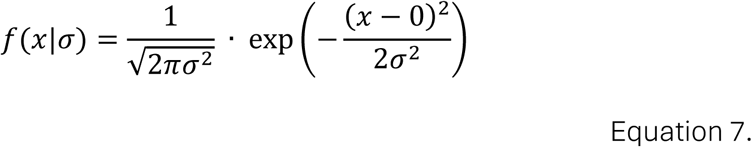

Where the standard deviation *σ* of the distribution in Equation 7 is:

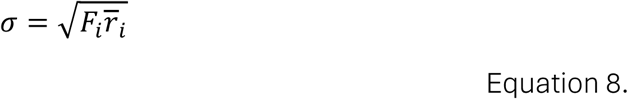

Noisy firing rates had a variance proportional to the mean firing rate, approximating a Poisson-like process, that was scaled by the Fano factor *F*_i_ with a default value of 1. Firing rates were constrained to positive values only, such that [*r*_i_]_+_ = *max*(*r*_i_ = 0).

#### iii. Orientation anisotropies

Four independent orientation anisotropies (differences between neurons selective for cardinal and oblique orientations) were implemented in the model: number of neurons *N*, orientation bandwidth ℎ, response variability *F*, and response gain *g* (Figure 2). Parameter values varied as a function of orientation according to a cosine function, with maxima and minima aligned to cardinal (0 and 90 °) and oblique (45 and 135 °) orientations. To control the magnitude of anisotropies in bandwidth, variability, and gain, the oblique parameter was held constant while the corresponding cardinal parameter was scaled to achieve the desired ratio.

**Figure 2.**
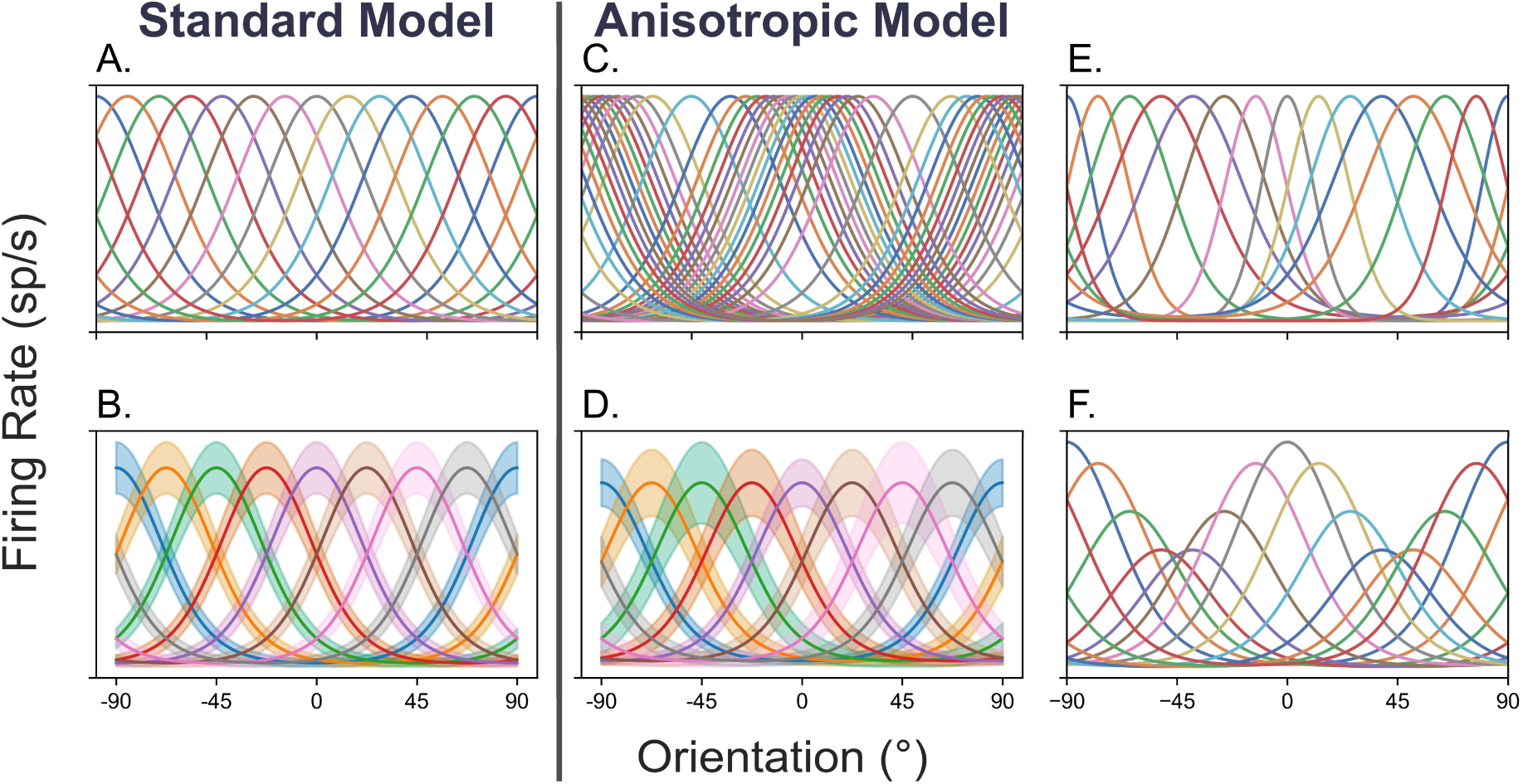
Visualisation of the model architecture. A reduced number of neurons is shown for visual clarity. (A) Orientation tuning curves for each neuron are defined by its preferred orientation, orientation bandwidth, and response gain. (B) Poisson-like noise is applied to neuronal responses on each trial. In the default model, all parameters are uniform across neurons (*left column*). Orientation anisotropies are introduced by varying parameter values as a function of orientation (*central and right columns*), including (C) the relative number of neurons preferring cardinal and oblique orientations, (D) response variability, (E) orientation bandwidth, and (F) response gain.

When quantifying the relative number of neurons preferring different orientations, cardinal-tuned neurons were defined as those with orientation preferences within ± 22.5 ° of 0 or 90 °, whereas oblique-tuned neurons had orientation preferences within ± 22.5 ° from 45 or 135 °. To adjust the relative number of cardinal and oblique neurons, orientation preferences were sampled from a cosine-shaped probability distribution, with the scale parameter optimised to produce the desired ratio.

#### iv. Spatial frequency

In macaque V1 neurons, the average orientation bandwidth decreases with increasing preferred spatial frequency [23,34]. The following equation describes this relationship for spatial frequencies above 1 c/° (adapted from Chen et al., 2020):

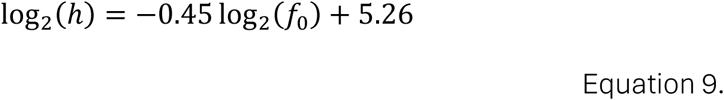

Where *f*_O_ is the preferred spatial frequency, and ℎ is the orientation bandwidth (HWHM). To simulate performance at the same spatial frequencies as the behavioural experiment (2, 8 and 16 c/°), independent model simulations were run with the respective default orientation bandwidths (28.1, 15.0 or 11.0 °) calculated from Equation 9.

Although fewer neurons prefer higher spatial frequencies than lower ones [51], simulations showed that population size did not substantially affect the magnitude of the oblique effect for either contrast detection or orientation discrimination (see supplementary materials, Figure S1 and S2). Thus, to simplify the interpretation of results, the number of neurons in the model was fixed (arbitrarily) at 540 at each spatial frequency.

#### v. Contrast detection

##### Pooling

Population responses *R* were calculated from the firing rates of neurons (Figure 3) according to the Minkowski method, where the contribution of each neuron is weighted by the magnitude of its response [52]:

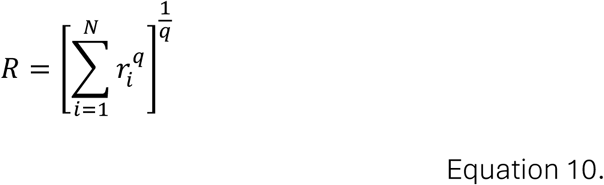

**Figure 3.**
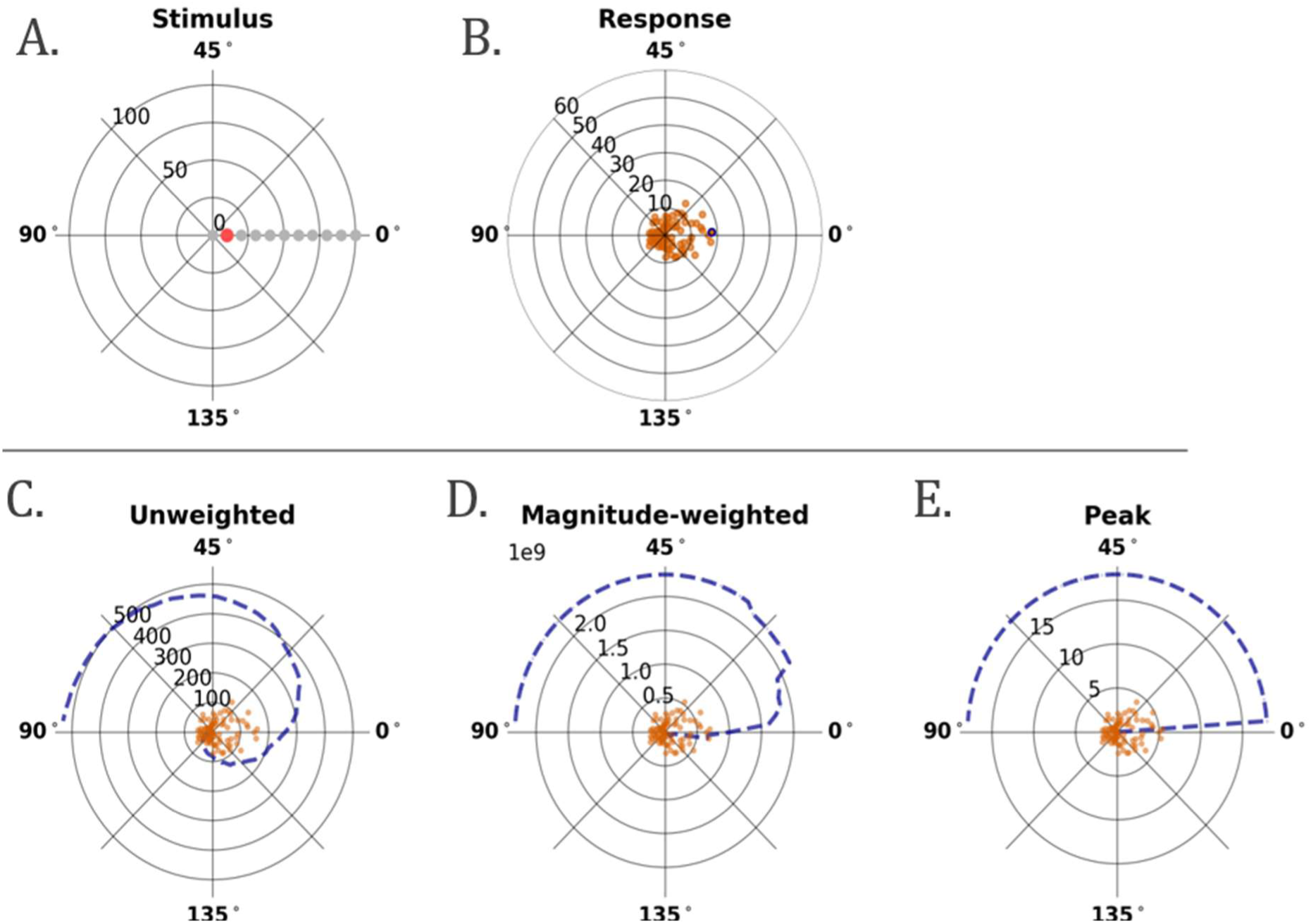
Visualisation of model responses (*top row*) and pooling methods (*bottom row*) using polar plots. (A-E) Polar angle represents orientation. (A) Stimuli are defined by orientation and contrast; contrast is represented by the radial distance from the origin. (B) Noisy firing rates in response to the stimulus, indicated by the red datapoint in panel A, are shown for neurons tuned to different preferred orientations. (C-E) Dotted lines show the cumulative sum of firing rates as responses are pooled over neurons with adjacent preferred orientations; radius represents the pooled response size, orange datapoints correspond to the firing rate values in panel B. (C) Unweighted pooling sums responses linearly from all neurons. (D) Magnitude-weighted pooling gives greater weight to larger responses. (E) Peak pooling selects only the strongest neuronal response. On each trial of the detection task, stimulus-present and stimulus-absent pooled responses are compared. Stimulus-absent responses are driven only by internal noise.

An exponent of *q* = 1 is used in the “unweighted” pooling condition, such that all neurons contribute equally to the population response *R*. An exponent *q* = 7 is used in the “magnitude-weighted” pooling condition. Alternatively, the population response in the “peak” condition is given by the firing rate of the maximally responding neuron, such that *R* = max *R*(*θ*).

##### Simulating performance

On each trial, two independent population responses were simulated for the standard orientation, at 0 % contrast (*c*_O_) and at a level of contrast (*c*_lvl_) determined by the staircase. The model decided which population response was larger:

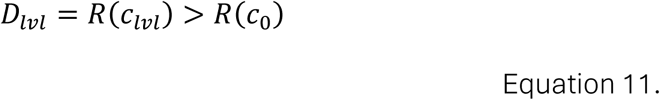

Where *D*_lvl_ is an array of binary values corresponding to the correctness of trial-to-trial responses at each level of contrast. The proportion of correct responses at each level was fitted with a logistic curve. Model thresholds were determined as the contrast value on the curve corresponding to 79.4 % correct responses, aligning with the point on the psychometric function targeted by the 3-down 1-up staircase procedure used in the behavioural experiment [53].

#### vi. Orientation discrimination

##### Orientation decoding

Stimulus orientation was decoded from the population firing rates by one of three commonly encountered decoding (“read-out”) algorithms (Figure 4). The winner-take-all (WTA) decoder is a heuristic method whereby the estimated orientation *θ*_WTA_ is equal to the orientation preference of the maximally responding neuron:

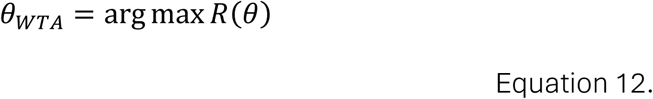

**Figure 4.**
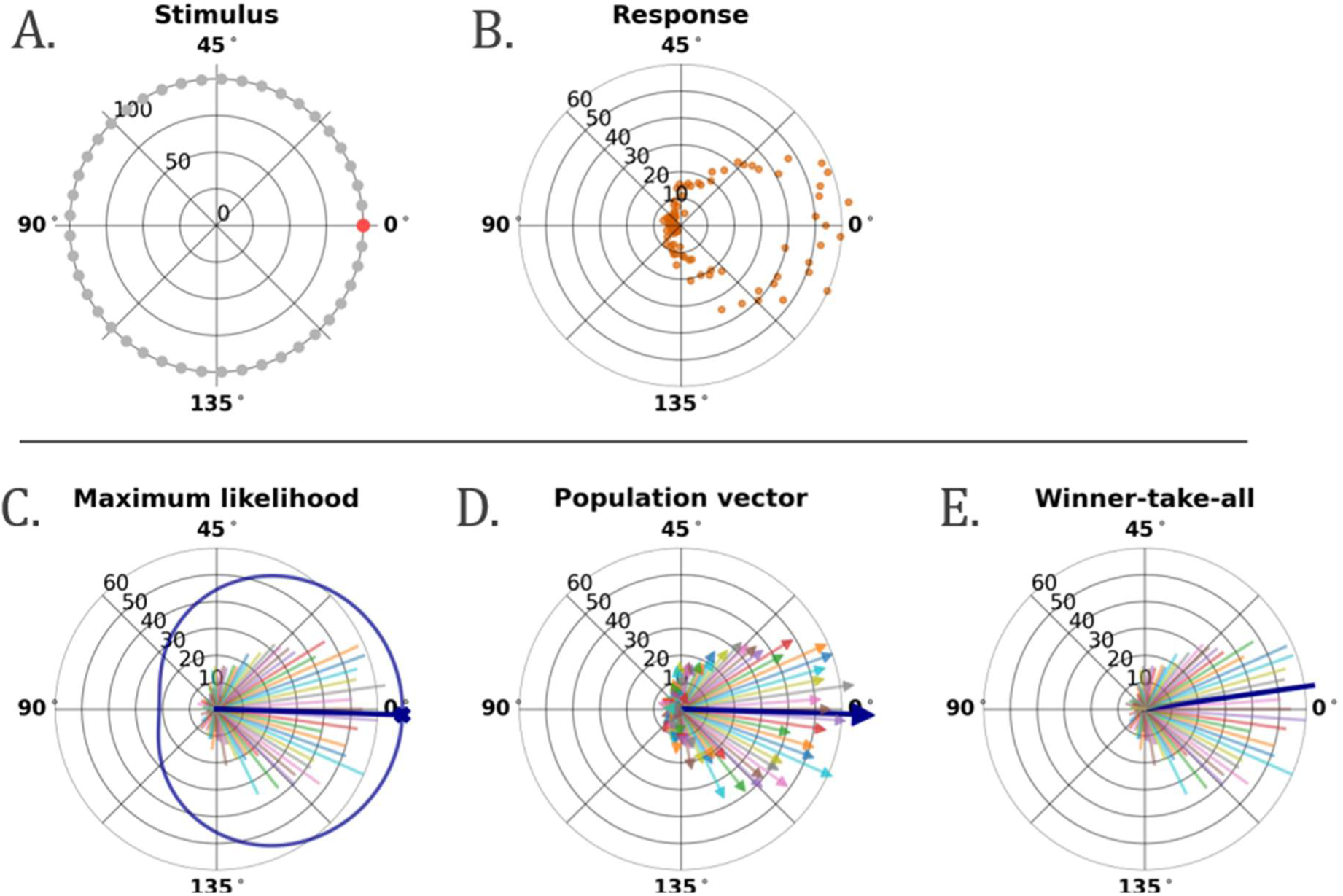
Visualisation of the model response (*top row*) and orientation decoders which estimated the stimulus orientation from the neuronal responses (*bottom row*). (A-E) Polar angle represents orientation. (A) Stimuli are defined by orientation and a fixed contrast of 100%; contrast is represented by radial distance from the origin. (B-E) Radius represents the neuronal firing rate. (B) Noisy firing rates in response to the stimulus, indicated by the red datapoint in panel A, are shown for neurons tuned to different preferred orientations. (C-E) Dark blue lines show the decoded orientation, estimated from the firing rates of neurons with different preferred orientations. (C) The maximum likelihood decoder selects the orientation with the highest likelihood, calculated from the orientation tuning curves and observed responses of all neurons. (D) The population vector decoder estimates orientation as the mean orientation vector, where each neuron’s response is represented by a vector defined by its preferred orientation and response magnitude. (E) The winner-take-all decoder estimates orientation as the preferred orientation of the neuron with the strongest response. On each trial of the discrimination task, decoded orientations are compared between two stimuli with slightly different orientations.

The population vector (PV) estimate *θ*_PV_ is the vector sum of all neuronal responses:

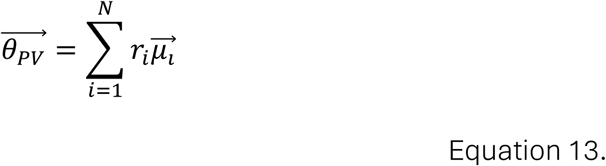

The maximum likelihood (ML) decoder estimates orientation from the weighted sum of neuronal responses by multiplying the firing rate of each neuron by the log of its tuning function [54]:

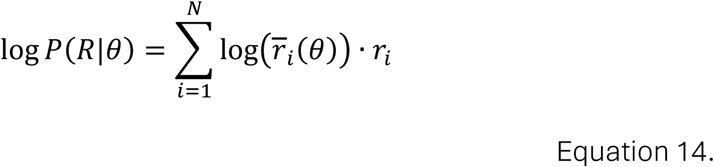

Such that the maximum likelihood estimate *θ*_ML_ is the orientation *θ* that maximises *P*(*R*|*θ*):

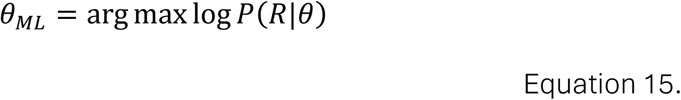

##### Simulating performance

The model decoded estimates of the standard *θ^*_std_ and comparison *θ^*_comp_ orientations at various levels of orientation difference, and then decided which estimate was more clockwise:

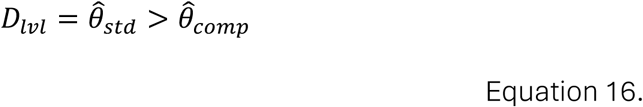

Model orientation-discrimination thresholds were calculated as described for contrast detection.

#### vii. Parameter estimation

Anisotropy values for number, bandwidth, response gain, and response variability were estimated by minimising the mean-squared error (MSE) between model and behavioural OEI scores. To avoid implausible solutions, cardinal and oblique parameter values were constrained to lie between a ratio of 1:4 and 4:1. Parameter values were estimated using a global optimisation algorithm, differential evolution (DE), due to its robustness to stochastic variability in the model output [55,56]. Differential evolution was run three times, using a population size of 12 candidates per free parameter over 10 iterations. During optimisation, candidate parameter sets evolve through selection and recombination, progressively concentrating sampling in regions of low error. The 5 % of parameter sets with the lowest error across all model iterations (at least 1200 fits per model) were then re-evaluated five times to obtain stable error estimates and were used to compute the median and interquartile range of each optimised free parameter.

## Results

### Experiment 1: Contrast detection

#### Behavioural results

At 2 c/°, there was little difference between cardinal (0 and 90 °) and oblique (45 and 135 °) contrast-detection thresholds, for all eight participants (Figure 5A and 5B). However, the size of the oblique effect increased with spatial frequency, with a geometric-mean OEI (± SE) score of 1.06 (± 0.04) at 2 c/°, increasing to 1.39 (± 0.21) at 8 c/°, and to 1.72 (± 0.19) at 16 c/° (Figure 5C). At 16 c/°, there were no systematic differences between thresholds for the left and right oblique, whereas seven out of eight participants showed lower thresholds for horizontal (0 °) relative to vertical (90 °). Cardinal and oblique thresholds were approximately 10 and 16 times higher at 16 c/° than at 2 c/°, respectively.

**Figure 5.**
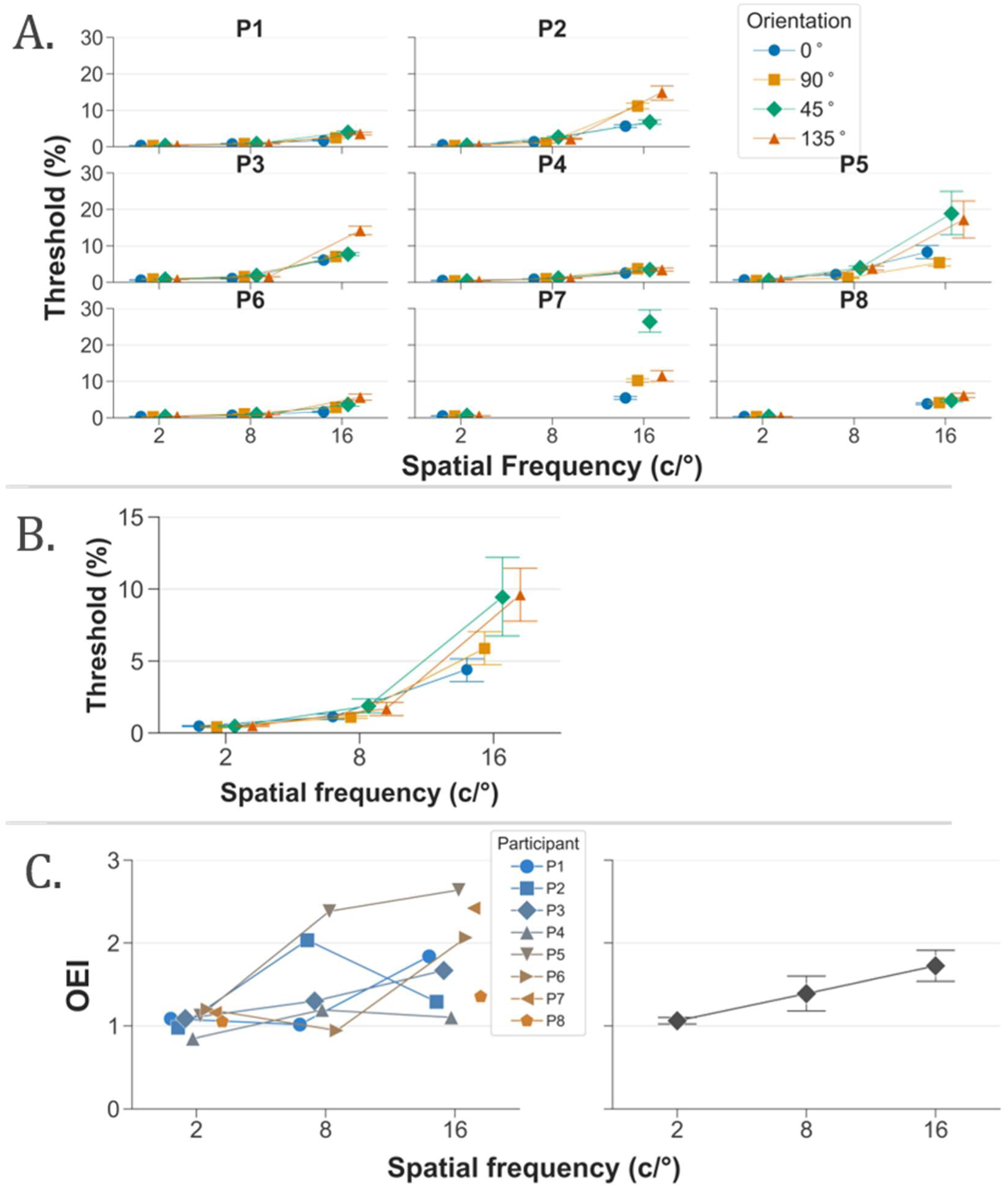
Behavioural results of Experiment 1. Individual (A) and mean (B) contrast-detection thresholds with error bars showing the standard error. (C) Individual and geometric mean oblique effect index (OEI) scores show the size of oblique effect. Adjoining lines are omitted for two participants (P7 and P8) who did not complete the 8 c/° condition. Oblique thresholds are similar to cardinal thresholds at 2 c/° but become larger than cardinal thresholds at 8 and 16 c/°.

#### Simulating the oblique effect for contrast detection

To examine whether combinations of anisotropies could account for the average behavioural oblique effect in contrast detection, anisotropies in number, bandwidth, response gain, and response variability were optimised together (Figure 6A and 6B). Various combinations of parameters produced the best-fitting data for each of the unweighted (UW), magnitude-weighted (MW) and peak (PK) pooling methods. However, all fits showed a cardinal overrepresentation (*N*_C_⁄*N*_O_ > 1.5) and slightly broadened cardinal bandwidths (ℎ_C_⁄ℎ_O_ > 1.3). Cardinal biases in response gain and response variability were less consistent across pooling methods. All pooling methods produced OEI scores consistent with the average behavioural data.

**Figure 6.**
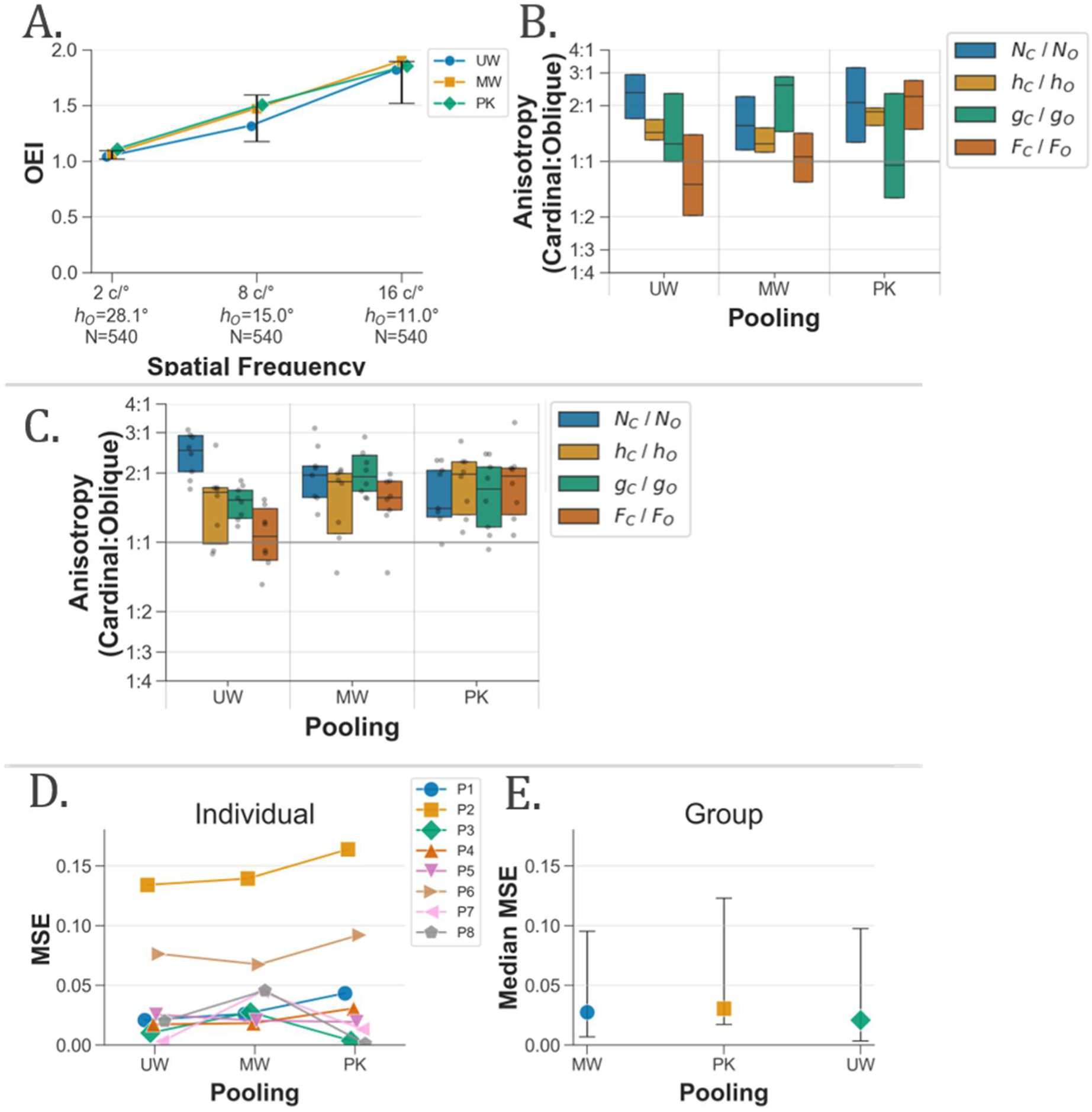
(A) Best-fitting oblique effect index (OEI) scores for contrast detection were simulated with anisotropies in number (*N*), bandwidth (*h*), gain (*g*), and noise (*F*) as free parameters. Error bars represent the standard error of behavioural OEI scores. Coloured lines indicate whether the pooling method was unweighted *UW*, magnitude-weighted *MW* or peak *PK*. With optimised parameters, all pooling methods were consistent with the average behavioural OEI scores. (B) Coloured box plots show the interquartile range and median value of each free parameter for each pooling method. (C) Best-fitting anisotropy values were calculated separately for each participant (grey datapoints), with the majority showing a cardinal bias in number, bandwidth, and gain. (D) Mean-squared error of model fits for each participant and (E) the median error across participants were similar for all pooling methods.

To evaluate which anisotropies systematically explained the behavioural data, rather than resulting from overfitting to the average data, best-fitting parameters were calculated for each individual participant (Figure 6C), and the fit error was compared for each pooling method (Figure 6D and 6E). Consistent with fits to the average data, a cardinal overrepresentation and broad cardinal bandwidths were found in the majority of participants, although heightened cardinal response gain was also found for all pooling methods (*g*_C_⁄*g*_O_ > 1.5). Each of these anisotropies increase the amplitude of pooled responses to cardinal orientations relative to obliques. In general, there were no substantial differences in the median fit error for each pooling method. Models with more restricted combinations of free parameters achieved similarly low fit errors for both the average and individual-participant data, for all pooling methods (see supplementary materials, Figure S3 and S4).

Spatial frequency was manipulated in the model through progressively narrower orientation bandwidths at higher spatial frequencies. At 2 c/°, the simulated oblique effect was either small or absent, demonstrating how broad bandwidths promote orientation-invariant detection performance by limiting the influence of anisotropic parameters in the neural population. It follows that the emergence of the oblique effect at higher frequencies is attributable to the uniform narrowing of orientation bandwidths. The detection of cardinal orientations is effectively prioritised when overall visual sensitivity is reduced at high spatial frequencies. However, the scale of loss in visual sensitivity at high spatial frequencies was not replicated by any best-fitting model (see supplementary materials, Figure S5).

### Experiment 2: Orientation discrimination

#### Behavioural results

Orientation-discrimination thresholds were substantially better for the cardinals than the obliques in each spatial frequency condition, consistent across all eight participants (Figure 7A and 7B). Geometric-mean OEI (± SE) scores increased with increasing spatial frequency, from 2. 48 (± 0.21) at 2 c/°, to 3.25 (± 0.29) and 3.36 (± 0.55) at 8 and 16 c/° (Figure 7C), however, in two participants (P2 and P5), smaller OEI scores were found at 16 c/° than 2 c/°. There was little consistency across participants in how cardinal thresholds changed with spatial frequency, whereas oblique thresholds at 16 c/° were higher than at 2 c/° for all eight participants. There was no general pattern in the differences between thresholds for vertical and horizontal, nor between the left and right obliques. Stimuli were presented at a fixed multiple of each participant’s contrast-detection threshold in each orientation and spatial frequency condition. Therefore, results cannot be attributed to gross variations in stimulus visibility.

**Figure 7.**
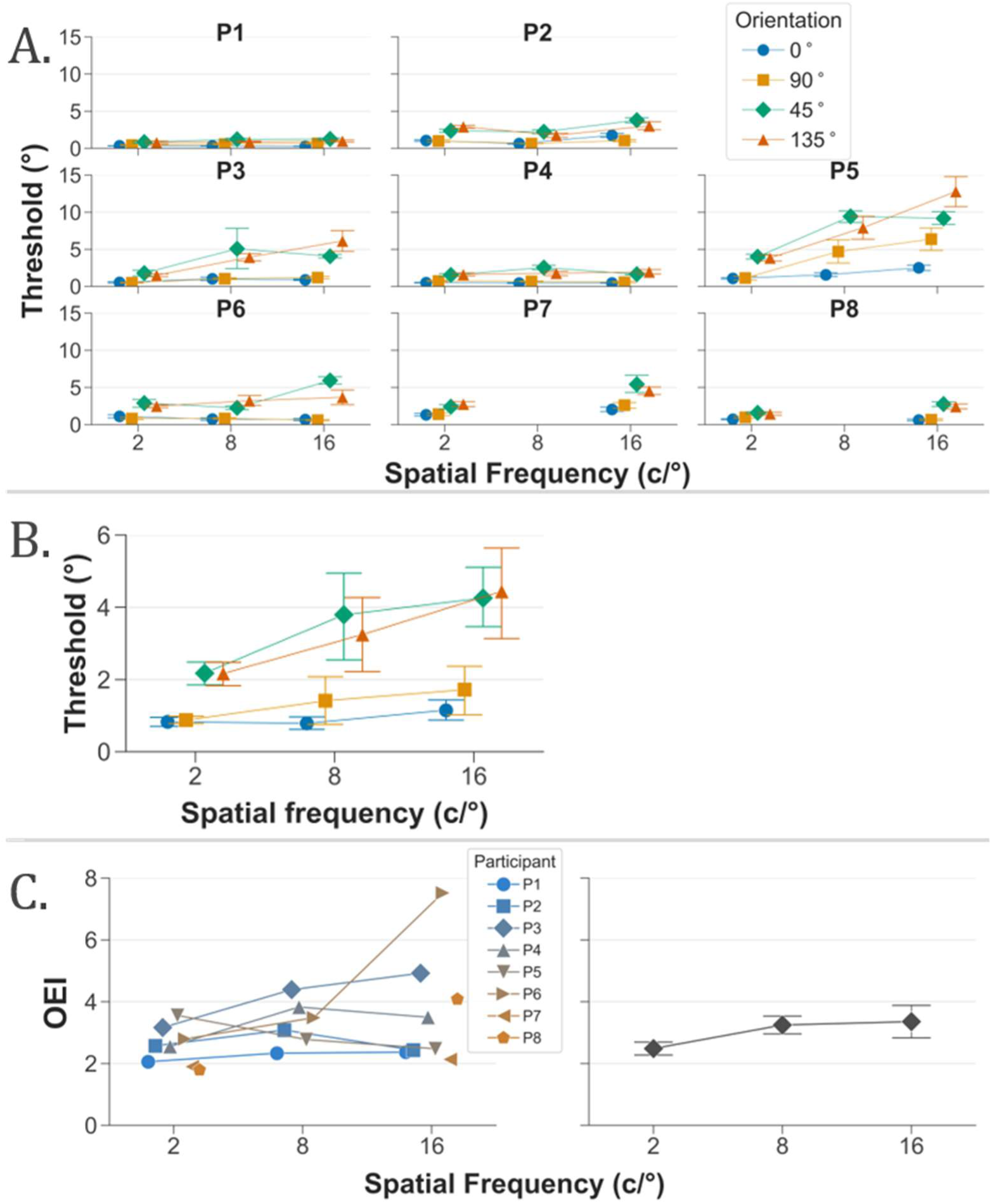
Behavioural results of Experiment 2. Individual (*A*) and mean (*B*) orientationdiscrimination thresholds with error bars showing the standard error. (C) Individual and geometric mean oblique effect index (OEI) scores for orientation discrimination. Adjoining lines are omitted used for two participants (P7 and P8) who did not complete the 8 c/° condition. Oblique thresholds are higher than cardinal thresholds at all spatial frequencies.

#### Simulating the oblique effect for orientation discrimination

To examine whether combinations of anisotropies could capture the average oblique effect in orientation discrimination, anisotropies in number, bandwidth, response gain, and response variability were optimised together (Figure 8A and 8B). Anisotropic bandwidths (ℎ_C_⁄ℎ_O_ < 0.35) featured in all model fits, alongside a cardinal overrepresentation (*N*_C_⁄*N*_O_ > 1.8) in ML and PV models, or heightened cardinal response gain for the WTA model (*g*_C_⁄*g*_O_ = 1.3). Only the WTA model produced OEI scores consistent with the behavioural data at all spatial frequencies. However, additional analyses revealed that WTA discrimination thresholds decreased greatly between 2 and 8 c/°, in stark contrast to the average behavioural data (see supplementary materials, Figure S5).

**Figure 8.**
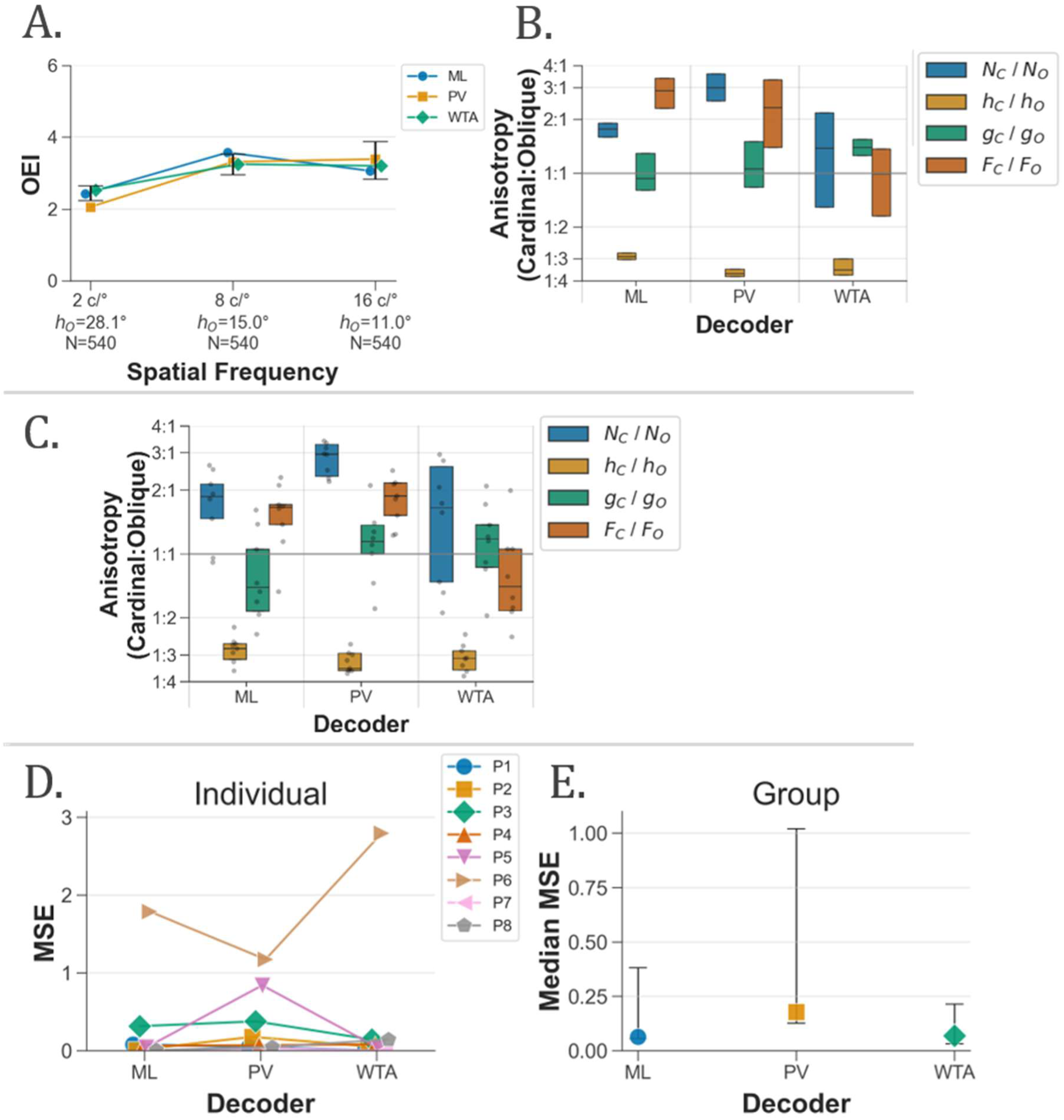
(A) Best-fitting oblique effect index (OEI) scores for orientation discrimination were simulated with anisotropies in number (*N*), bandwidth (*h*), gain (*g*), and noise (*F*). Error bars represent the standard error of behavioural OEI scores. Coloured lines indicate whether the orientation decoder was maximum likelihood *ML*, population vector *PV* or winner-take-all *WTA*. With optimised parameters, all decoders were generally consistent with the average behavioural OEI scores. (B) Coloured box plots show the interquartile range and median anisotropy value of each free parameter for each orientation decoder. (C) Best-fitting anisotropy values were calculated separately for each participant (grey datapoints), with all showing greatly narrowed cardinal bandwidths. (D) Mean-squared error of model fits for each participant and (E) the median error across participants were similar for ML and WTA decoders, whereas the PV decoder struggled to capture individual differences.

The best-fitting anisotropies for individual participants were generally consistent with the average data (Figure 8C), whereby cardinal bandwidths were much narrower than oblique bandwidths. However, curious features emerged for certain models, such as lowered cardinal response gain for the ML decoder, and heightened cardinal response variability for the ML and PV decoders. The median fit error across participants was lowest for the ML and WTA decoders, whereas the PV model struggled to capture individual differences in performance (Figure 4D and 4E). Low error was only found when bandwidth was a free parameter in the model fit (see supplementary materials, Figure S3 and S4). Notably, a cardinal overrepresentation was found in the majority of participants for all decoders, which is a key feature potentially shared between the models for contrast detection and orientation discrimination.

#### Shared model of the oblique effect for detection and discrimination

To formally test whether a shared model could account for the oblique effects in contrast detection and orientation discrimination, a final series of simulations fit all anisotropies to the average behavioural OEI scores of both tasks simultaneously (Figure 9). A singular model could not replicate the patterns of results found for the behavioural oblique effect on both tasks, not for any combination of detection pooling methods and orientation decoders. Greatly narrowed cardinal bandwidths were required to capture the behavioural oblique effect in orientation discrimination, but their inclusion directly prevented the model from replicating the detection oblique effect. Shared models failed to capture the absence of an oblique effect at 2 c/° on the detection task, and the increasing OEI scores between 2, 8 and 16 c/°. These findings indicate that the oblique effects on both tasks are driven by distinct orientation biases in independent neural populations.

**Figure 9.**
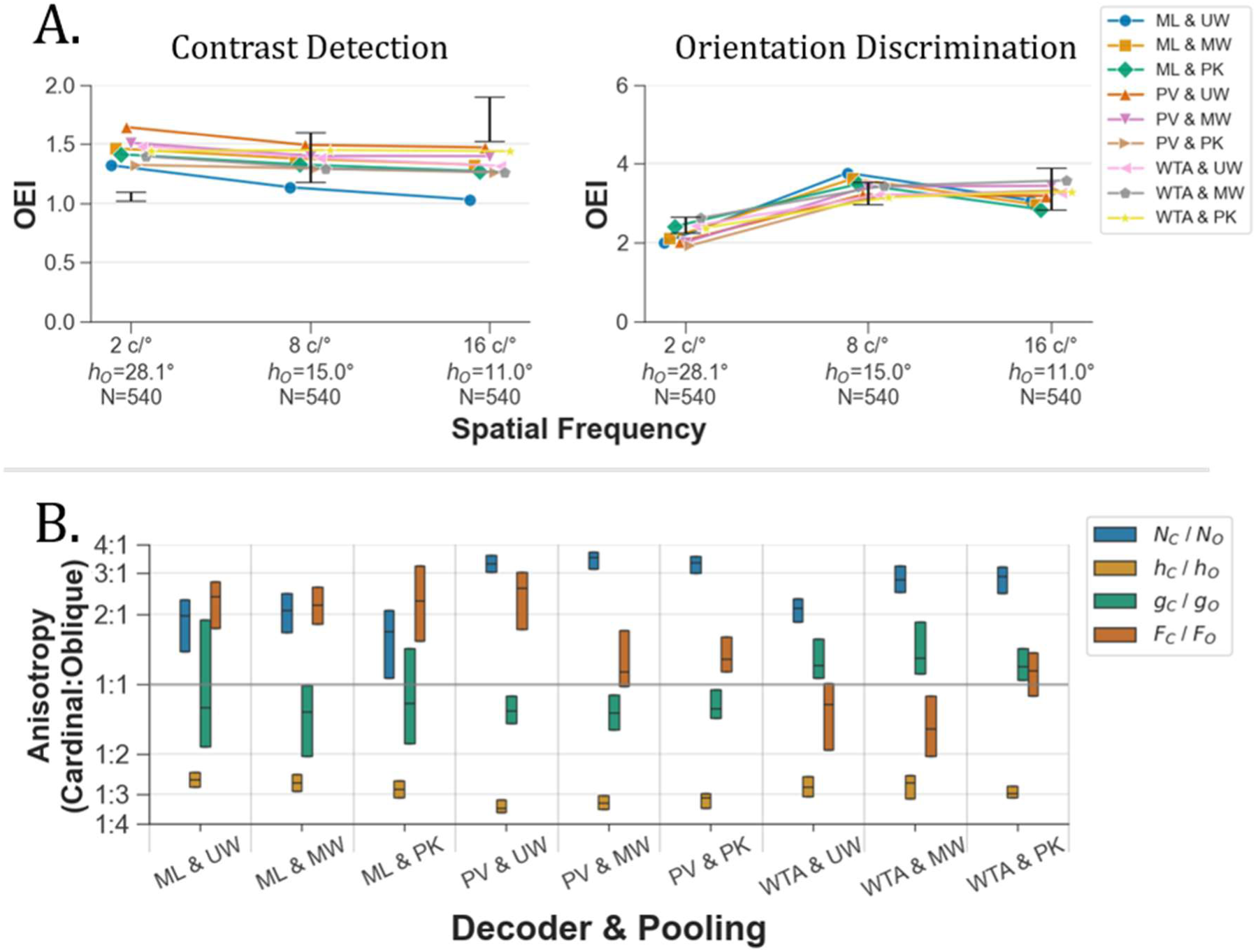
(A) Best-fitting oblique effect index (OEI) scores for model fits shared across both the contrast-detection and orientation-discrimination tasks, with anisotropies in number (*N*), bandwidth (*h*), gain (*g*), and noise (*F*) as free parameters. Error bars represent the standard error of the behavioural OEI scores. (B) Coloured box plots show the interquartile range and median anisotropy value of the best-fitting parameters for each combination of orientation decoder and detection pooling method. The detection pooling method was either unweighted *UW*, magnitude-weighted *MW* or peak *PK*, and the discrimination orientation decoder was either maximum likelihood *ML*, population vector *PV*, or winner-take-all *WTA*. Results show that a shared model could not explain the behavioural oblique effect in both tasks simultaneously.

## Discussion

To investigate whether orientation encoding is determined by a fixed strategy or is flexible, dependent on task demands, we provide novel computational accounts of the oblique effects for contrast detection and orientation discrimination. We adjusted parameters of a neurophysiologically-inspired model of orientation encoding to account for behavioural performance on both tasks. By fitting the model to the data of individual participants, we identified which variables systematically explained the behavioural data rather than resulting from overfitting to the average data. A general finding was that bandwidths of orientation-selective neurons, known to uniformly decrease with increasing spatial frequency in macaque V1 [23,34], can significantly modulate the size of the oblique effect.

Consistent with classic studies [6,7], the behavioural oblique effect for contrast detection was absent at 2 c/° but increased robustly with increasing spatial frequency. The absence of oblique effect at lower spatial frequencies in the model was attributable to the broad, overlapping orientation-tuning curves of responding neurons (including those with orthogonal orientation preferences), such that the responses of cardinal-tuned neurons facilitated the detection of both cardinal and oblique orientations. At higher spatial frequencies, the extent of neuronal recruitment for a given stimulus orientation is reduced due to narrowed orientation bandwidths, meaning that greater responses from cardinal-tuned neurons can specifically enhance perception of cardinal versus oblique orientations. This enhancement comes at the cost of impaired detection of oblique orientations due to the influence of spontaneous firing (noise) of unrecruited cardinal-tuned neurons. Our account is consistent with reports of a cardinal overrepresentation in V1 of humans [14], V1/V4 in macaques [15], and striate cortex in cats [11], found at spatial frequencies well-below those that elicit the oblique effect for contrast detection. An oblique effect in contrast detection at relatively high spatial frequencies may serve to offset marked losses in overall sensitivity, favouring cardinal orientations which are more prevalent in typical visual experience [3–5] across all spatial scales [35].

As expected from previous work, the oblique effect for orientation discrimination was present at all tested spatial frequencies, and oblique thresholds increased on average with increasing spatial frequency [36,37]. Anisotropic orientation bandwidths, narrower for cardinal-tuned than oblique-tuned neurons, were necessary to simulate the oblique effect in orientation discrimination. This was the case for each of the maximum likelihood (ML), population vector (PV), and winner-take-all (WTA) decoders. Additional anisotropies in neuronal number, response gain, and response variability enabled the model to more accurately simulate the behavioural oblique effect. However, the PV model struggled to account for individual differences in behavioural performance. To further explore which decoding methods are employed during these tasks, future research (in progress) should test predictions of the WTA, PV, and ML models under different experimental conditions. For example, stimuli could be presented at near-threshold contrasts or under varying levels of external noise, to explore whether flexible orientation read-out algorithms are employed for stimuli of differing visibility [38,39].

A shared model could not simultaneously account for the behavioural oblique effects in both contrast-detection and orientation-discrimination tasks, regardless of the combination of pooling method and orientation decoder. Anisotropic orientation bandwidths were necessary to simulate the oblique effect for orientation discrimination, but their inclusion directly prevented capturing the oblique effect for contrast detection. Assuming that the oblique effect reflects a functional adaptation to the prominence of cardinal orientations in the visual environment [3–5], whereby accurate perception of cardinal orientations is prioritised over oblique orientations, our findings suggest that such adaptations may occur in independent subpopulations of neurons that serve particular visual functions. For example, discrimination may rely on neural circuits with narrowed cardinal orientation bandwidths, whereas detection may utilise an independent neural population with uniform bandwidths and heightened responses of cardinal-tuned neurons. Such an arrangement would prioritise cardinal information for one task while leaving representations for other tasks unaffected.

These ideas align with the typical pattern of perceptual learning, where improvements rarely generalise across different visual tasks [40], even when training induces measurable changes in visual cortex [41,42]. Indeed, visual experience leads to neural adaptations which are tightly tuned to the demands of the trained task [25,27,28]. For example, orientation-discrimination training led to narrowed orientation bandwidths in monkey V4 neurons, which was most pronounced in neurons with steeper slopes, i.e., whose tuning was most informative for signalling differences at the trained orientation [27,28]. On the other hand, training on a detection task at low-to-mid spatial frequencies led to enhanced neuronal contrast gain in cat area 17 [24], whereas training at high spatial frequencies resulted in a reconfigured distribution of preferred spatial frequencies [25]. Importantly, the reason why single-cell studies in macaque V1 and V2 often report no cardinal biases [22,23] may be because such biases are expressed in independent neural circuits supporting different perceptual judgements, rather than being general properties of visual cortex.

Our findings suggest that the brain deploys distinct computational strategies when encoding orientation for detection and discrimination tasks, with dissociable neural circuits which are independently shaped by environmental statistics and task demands. The flexibility of this framework serves to boost perceptual performance for cardinal orientations relative to oblique orientations across a variety of tasks. Indeed, a growing body of evidence indicates that neurons can dynamically adapt their response properties in order to prioritise behaviourally relevant information [43]. For example, single neurons in inferotemporal cortex exhibit pronounced context-dependent shifts in selectivity, transitioning from broadly tuned object detectors to highly selective face-responsive neurons depending on the stimulus category [44]. Taken together, these findings point to a more general principle, that sensory representations are not fixed, but flexible, and can be reconfigured to meet different behavioural demands.

## Acknowledgements

Code availability: The data analysis and modelling code is available from https://doi.org/10.5281/zenodo.20544481

## Declarations

The authors have no relevant financial or nonfinancial interests to disclose.

## Ethics approval

The experimental procedures were approved by the local ethics committee (School of Psychology, University of Nottingham, UK).

## Author Contributions

RL: Conceptualisation, Methodology, Software, Formal analysis, Data curation, Visualisation, Writing – original draft, Writing – review and editing.

TL: Conceptualisation, Methodology, Supervision, Project administration, Writing – review and editing.

PM: Conceptualisation, Methodology, Supervision, Project administration, Writing – review and editing.

